# Reversing myeloid-derived suppressor cells mediated immunosuppression via p38α inhibition enhances immunotherapy efficacy in triple negative breast cancer

**DOI:** 10.1101/2023.03.31.535102

**Authors:** Qianyu Wang, Shasha Li, Yifei Dai, Xiankuo Yu, Yumei Wang, Lu Li, Ming Yang, Kequan Lin, Wei Shao, Haiyan Wang, Huili Wang, Guanbin Zhang, Dong Wang

## Abstract

Infiltration of myeloid-derived suppressor cells (MDSCs) leads to Immunosuppressive tumor microenvironment (TME), which is one of the major causes for low objective response rates of immune checkpoint blockade (ICB) therapy. Here, we report that chemical inhibition of p38α reverses this MDSC-induced immunosuppressive TME and improves the immunotherapy efficacy in triple negative breast cancer (TNBC). Firstly, by combining the tumor immunological phenotype (TIP) gene signature and high throughput sequencing based high throughput screening (HTS^2^), we identified that ponatinib significantly inhibits the expression of “cold” tumor associated chemokines CXCL1 and CXCL2 in cancer cells. This inhibition decreases the infiltration of MDSCs and consequently increased the accumulation of “hot” tumor associated T cells and NK cells and thus reverses the immunosuppressive TME. Then, by multiple preclinical models, we found that ponatinib significantly inhibits tumor growth in a TME-dependent manner and enhances the efficacy of anti-PD-L1 immunotherapy on TNBC *in vivo*. Notably, ponatinib exhibits no significant inhibition on immune cells in mouse spleens. Mechanistically, ponatinib directly inhibits the kinase activity of p38α, which results in the reduction of the phosphorylation of STAT1 at Ser727, and thus the decreased expression of CXCL1 and CXCL2 in cancer cells. Our study provided the therapeutic potential of combining p38α inhibition with ICB for the treatment of TNBC.

## Introduction

Cancer immunotherapies, which modulate the body’s own immune system to against tumor growth, have become effective therapeutic strategies for cancer treatment (Hamid et al., 2013). Recently, immune checkpoint blockade (ICB) therapy that targets cytotoxic T lymphocyte-associated protein 4 (CTLA-4) or programmed cell death protein 1/ programmed cell death ligand 1 (PD-1/PD-L1) has been widely proved to have remarkable antitumor responses (Brahmer et al., 2012; Hamid et al., 2013). Unfortunately, only a small subset of patients receiving immunotherapy will respond to these immune checkpoint inhibitors (ICI) (Brahmer et al., 2012; Hamid et al., 2013). Growing evidence suggests that one reason for the low objective response rate and poor survival in solid tumors is closely related to the immunosuppressive TME (Bianchini et al., 2016). Therefore, it is expected that combination immunotherapy targeting the immunosuppressive TME will be needed to improve the efficacy and responsiveness of ICB therapy(Sharma et al., 2017).

Based on the degree of immune cell infiltration in the tumor microenvironment, tumors are now frequently classified as "cold" (non-inflamed) or "hot" (inflamed) phenotypes (Gajewski, 2015; Nagarsheth et al., 2017). The immune landscape of “cold” tumors is characterized by the immunosuppressive TME, such as predominantly myeloid cells infiltration and the lack of T cells. MDSCs and macrophage constitute the majority of infiltrating immunosuppressive cells, which accumulation negatively suppress T cell infiltration and function and have been widely investigated in various cancer types. It is shown that intratumoral MDSCs drive resistance to immunotherapy, thereby promoting tumor development and metastasis (Mauti et al., 2011; Veglia et al., 2018). Clinical researches have shown that MDSCs highly correlate with tumor burden and poor overall survival of various cancers including colorectal cancer, non-small cell lung cancer, liver cancer, melanoma, thyroid cancer and bladder cancerAngell et al. (2016); (Arihara et al., 2013; Diaz-Montero et al., 2009; Huang et al., 2013; Jordan et al., 2013; Li et al., 2017; Sun et al., 2012; Wang et al., 2018; Yang et al., 2017). Thereby, it indicates that MDSCs are potential therapeutic targets for cancer immunotherapy. Agents that can eliminate MDSCs may overcome resistance to ICB across cancer types.

MDSCs comprise two different subsets: polymorphonuclear (PMN-) MDSCs and monocytic (M-) MDSCs (Youn et al., 2008). Selective cytokines and tumor-associated factors such as CXCL1, CXCL2, IL6, IL10, TNF-α, GM-CSF, NF-κB, STAT, etc., drive the MDSCs development (Li et al., 2021). Currently, investigations targeting MDSCs have been focused on MDSCs recruitment. The migration of newly formed MDSCs into tumor microenvironment is mainly driven by C–X–C motif chemokine ligand 1 (CXCL1) and CXCL2 and their receptors CXCR1 and CXCR2. (Stadtmann and Zarbock, 2012; Steele et al., 2016). Therefore, an alternative approach to decrease numbers of tumor-infiltrating MDSCs is to interfere MDSCs recruitment, such as block CXCL1/2-CXCR1/2 axis. Additionally, foundational studies and our previous work have demonstrated that compound induced the expression of Th1 chemokines could convert immunologically "cold" tumors into "hot" tumors and thereby confers a therapeutic effect against cancer cells (Nagarsheth et al., 2016; Peng et al., 2015; Wang et al., 2021). Thus, combination therapy with small molecule that inhibit MDSCs infiltration into tumor microenvironment would be particularly attractive therapeutics for "cold" tumors.

The aim of the present study is to discover potential immunotherapy combination agents which could alter immune suppression. In this project, we revealed that the inhibition of p38α by ponatinib represses the expression of chemokine CXCL1 and CXCL2 in cancer cells via p38α-STAT1 signaling pathway, which reshapes the immune microenvironment by inhibiting MDSCs infiltration, thus restoring the antitumor immunity of T lymphocytes or NK cells. In addition, combining ICB agents and ponatinib synergistically suppressed growth of breast tumor *in vivo*. These findings provide a new combination immunotherapy to synergistically inhibit tumor growth.

## Results

### HTS^2^ screening identified that ponatinib inhibits the expression of CXCL1 and CXCL2 in TNBC cells

Our previous work has proved that using high-throughput drug screening for compounds identification is feasible (Shao et al., 2019; Wang et al., 2021). In consideration of the significant role of immunosuppressive TME, compounds targeting “cold” tumors need to be urgently discovered. In this project, to identify small-molecule compounds reshaped the immunosuppressive TME of “cold” tumors, a high-throughput drug screening containing 1,154 US Food and Drug Administration (FDA)-approved drugs treating with breast cancer cell line MDA-MB-231 for 24 h was performed (Shao et al., 2019).

The relative activity of each compound was scored according to its ability to reverse the expression of the selected TIP signature genes: “cold” tumor–related TIP genes *CXCL1* and *CXCL2*, and “hot” tumor–related TIP genes *CXCL10* and *CXCL11*. We used two connectivity methods (Lamb et al., 2006; Subramanian et al., 2017) to score and rank the screening compounds. Meanwhile, in our pursuit of discovering compounds that inhibit the expression of "cold" tumor-related TIP genes, we screen for compounds that result in a fold change (FC) of *CXCL1* and *CXCL2* expression less than −1 (log2FC<-1). Four compounds were identified that met all three criteria, which are Candesartan cilexetil (Atacand), Ponatinib (AP24534), Estriol and Ciprofloxacin (Cipro) (Fig.1A-1C). Notably, Candesartan cilexetil is an angiotesin-II receptor antagonist, a prodrug that is rapidly converted to candesartan, which can inhibit the expression of *CXCL1* induced by stroke (Schmerbach et al., 2008; Wang et al., 2018). This result demonstrated that our approach can effectively identify potential drugs which alter “cold” tumor–related TIP signature.

**Fig1.**
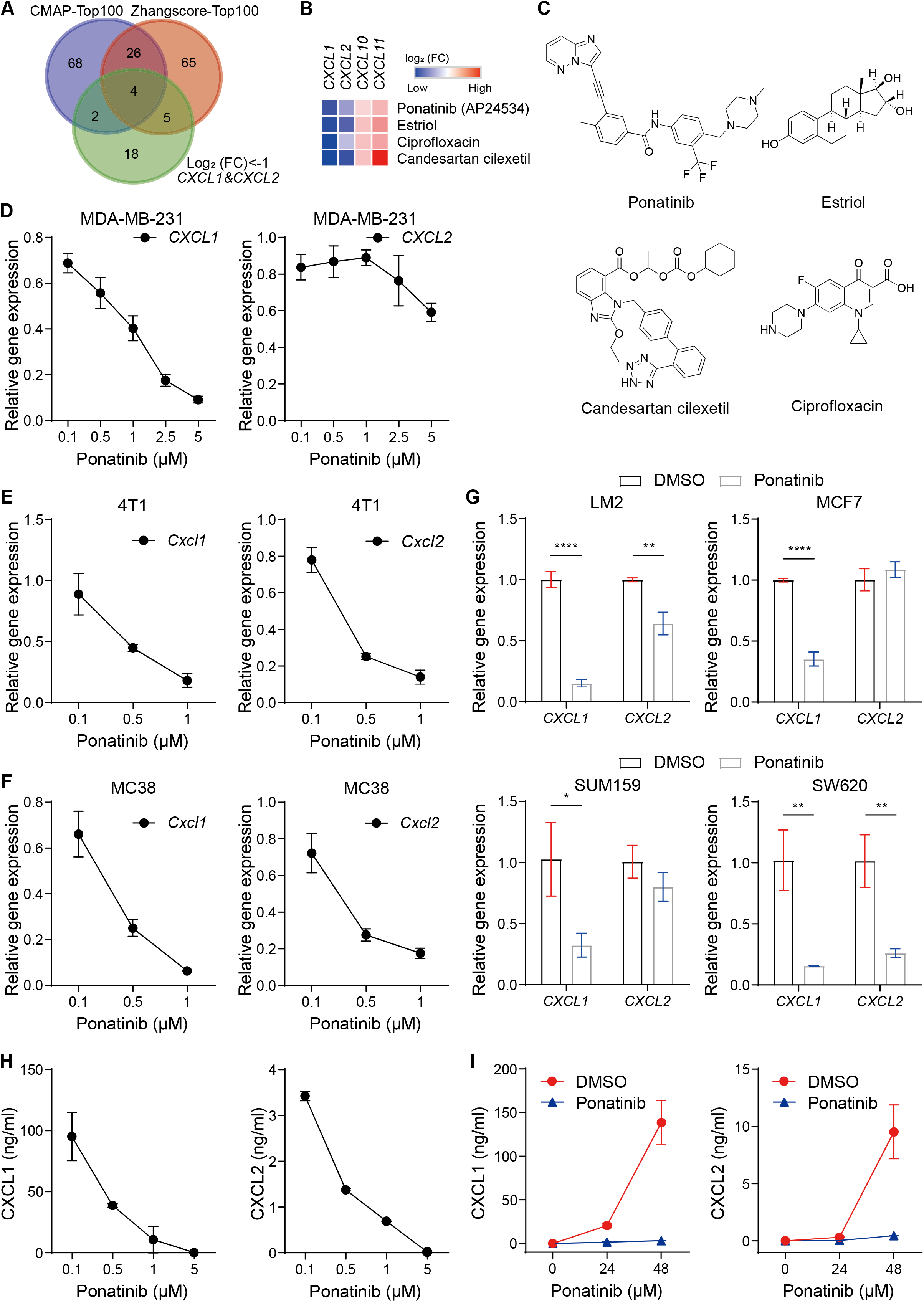
Ponatinib inhibits *CXCL1* and *CXCL2* expression in multiple cell lines. (A)Venn diagram analysis across three groups of drugs containing (i) top 100 drugs ranking by CMAP score; (ii) top 100 drugs ranking by zhangscore; (iii) fold change (*CXCL1* and *CXCL2*) >2. (B) Heat map representing the expression of indicated chemokine genes in MDA-MB-231 cells treated with the top 4 hit compounds, the log_2_ (fold change) is used to scale the differences in gene expression (red for up-regulation and blue for down-regulation) (C) The top 4 chemical structures are shown. (D, E and F) RT-qPCR analysis showed that ponatinib was capable of reversing the expression of the indicated chemokine genes in a dose manner. MDA-MB-231(D), 4T1 (E) and MC38 (F) cells were treated with different concentrations of ponatinib at 24 h. (G) RT-qPCR analysis of the indicated cell lines treated with the 1 μmol/L ponatinib at 24 h. (H and I) ELISA analysis of CXCL1 and CXCL2 in 4T1 cells treated with different concentrations of ponatinib (H) or with 1 μmol/L ponatinib at various time points (I). Data represent mean ± SD of three independent experiments and normalized to DMSO control.

Ponatinib, an FDA-approved drug for leukemia, has not been previously linked to reshape “cold” tumor related TME. Thus, we selected ponatinib for further investigation. To validate the expression changes of *CXCL1* and *CXCL2* at the mRNA levels after ponatinib treatment, quantitative reverse transcription polymerase chain reaction (RT-qPCR) was performed in multiple cancer cell lines. Our results confirmed that ponatinib indeed dose-dependently affected the mRNA expression levels of *CXCL1* and *CXCL2* in MDA-MB-231 cells (Fig. 1D). These genes exhibited similar expression pattern in murine breast cancer 4T1 cells, mouse colorectal cancer MC38 cells and other human breast cancer cells or colorectal cancer cells when treated with ponatinib (Fig. 1E-1G). Consistent with the effect on transcription level, enzyme-linked immunosorbent assay (ELISA) of cell exudates showed that ponatinib treatment also caused a significant decrease in secretion of both CXCL1 and CXCL2 chemokines from cancer cells (Fig. 1H-1I). Our findings suggested that ponatinib inhibits the expression of CXCL1 and CXCL2. Therefore, we selected ponatinib, identified through HTS2 screening, as a potential drug to target "cold" tumors for further investigation.

### Ponatinib inhibits TNBC growth in immunocompetent mice but not immune-deficient mice

To determine the impact of various tumor microenvironments on the efficacy of ponatinib in breast cancer *in vivo*, both immunocompetent and immune-deficient mammary tumor models were utilized. Firstly, the anti-tumor efficacy of ponatinib was compared in 4T1 transplantation in WT BALB/c mice (immunocompetent mice), nude BALB/c mice (immuno-deficient mice, lack T cells) and NSG mice (immuno-deficient mice, lack T cells, NK cells and B cells). Following intramammary infusion, 4T1 tumors in BALB/c nude mice and NSG mice were colonized to a similar extent to tumors of wild-type mice. As expected, after ponatinib treatment (Fig. 2A), both tumor volume and tumor weight were significantly suppressed in WT mice (66.2% and 73.3% inhibition ratio, respectively, *P<0.001*)(Fig. 2B) and nude mice (52.0% and 69.0% inhibition ratio, respectively, *P<0.0001*) (Fig. 2C). Notably, in the NSG mice, the same course of ponatinib treatment showed no significant differences on tumor volume and tumor weight (Fig. 2D).

**Fig2.**
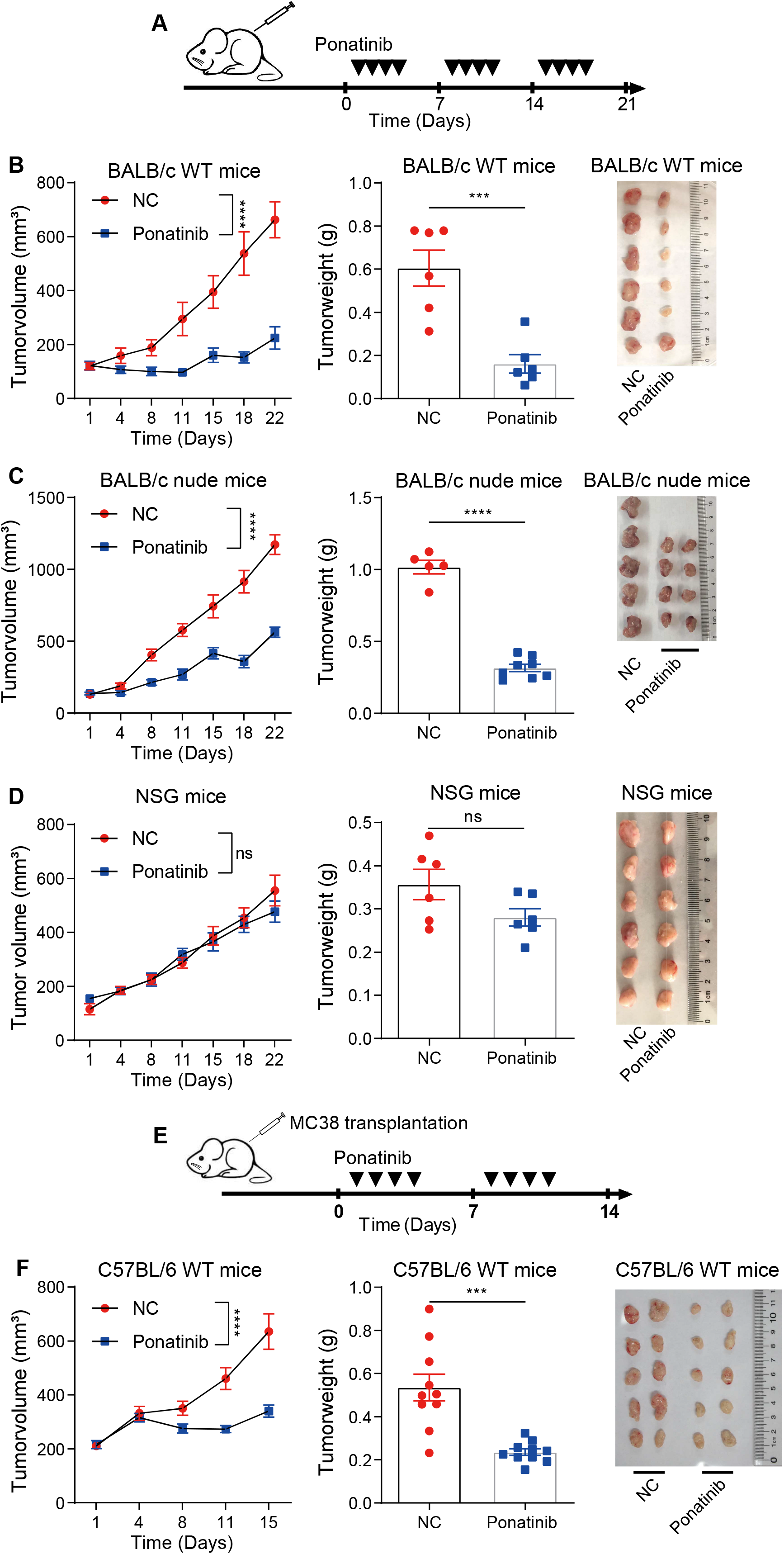
The anti-tumor effects of ponatinib depend on tumor microenvironments. (A) Schematic diagram for the *in vivo* drug studies in 4T1-bearing mice. Ponatinib (30mg/kg) or vehicle was administered four days one week when tumor volume reached 100 mm³. (B) Growth of 4T1 cells in BALB/c WT mice (left, n = 6 mice per group) treated with ponatinib or vehicle. Statistical significance was determined by two-way ANOVA. \*\*\*\**P* < 0.0001. Tumor weight (middle) was measured after the tumors were harvested of the indicated group. Statistical significance was determined by unpaired two-tailed Student’s t tests. \*\*\**P* < 0.001. Data represent mean ± SEM of n mice, as indicated on each panel. Images (right) of 4T1 tumors of the indicated groups. (C) Growth of 4T1 cells in BALB/c nude mice (left, control n = 5 mice; ponatinib = 8 mice) treated with ponatinib or vehicle. Statistical significance was determined by two-way ANOVA. \*\*\*\**P* < 0.0001. Tumor weight (middle) was measured after the tumors were harvested of the indicated group. Statistical significance was determined by unpaired two-tailed Student’s t tests. \*\*\*\**P* < 0.0001. Data represent mean ± SEM of n mice, as indicated on each panel. Images (right) of 4T1 tumors of the indicated groups. (D) Growth of 4T1 cells in NSG mice (left, n = 6 mice per group) treated with ponatinib or vehicle. Statistical significance was determined by two-way ANOVA. ns, *P* > 0.05. Tumor weight (middle) was measured after the tumors were harvested of the indicated group. Statistical significance was determined by unpaired two-tailed Student’s t tests. ns, *P* > 0.05. Data represent mean ± SEM of n mice, as indicated on each panel. Images (right) of 4T1 tumors of the indicated groups. (E) Schematic diagram for the *in vivo* drug studies in MC38-bearing mice. Ponatinib (30mg/kg) or vehicle was administered four days one week when tumor volumn reached 200 mm³. (F) Growth of MC38 cells in C57BL/6 mice treated with ponatinib or vehicle. (left, n = 10 mice per group). Data represent mean ±SEM. Statistical significance was determined by two-way ANOVA. \*\*\*\**P* < 0.0001. Tumor weight (middle) was measured after the tumors were harvested of the indicated group. Data represent mean ± SEM. Statistical significance was determined by unpaired two-tailed Student’s t tests.\*\*\**P* < 0.001. Images (right) of MC38 tumors of the indicated groups.

To extend our analysis, we subcutaneously injected MC38 cells in WT C57BL/6 mice (Fig. 2E). Strong inhibition of tumor growth was also observed in MC38 subcutaneous tumors in C57BL/6 mice under ponatinib treatment for two weeks. Both tumor volume and tumor weight were significantly decreased (Fig. 2F). Together, our results clearly showed that the anti-cancer effect of ponatinib is TME-depend. Meanwhile, our data also suggested that T cell and NK cell in TME are essential for the anti-cancer function of ponatinib.

### Ponatinib inhibits the infiltration of MDSCs into TME *in vivo* by suppressing the expression of CXCL1 and CXCL2 in cancer cells

To determine the TME-dependent anti-cancer function of ponatinib, the changes in TME after ponatinib treatment were investigated. Firstly, our results showed that ponatinib inhibits the expression of *Cxcl1* and *Cxcl2* in 4T1 tumors (Fig. 3A), which is consistent with our results *in vitro* (Fig. 1D-1G). It is well known that CXCL1 and CXCL2 are the two major CXCR2 ligands in attracting immunosuppressive MDSCs to the TME (Stadtmann and Zarbock, 2012; Steele et al., 2016).

**Fig3.**
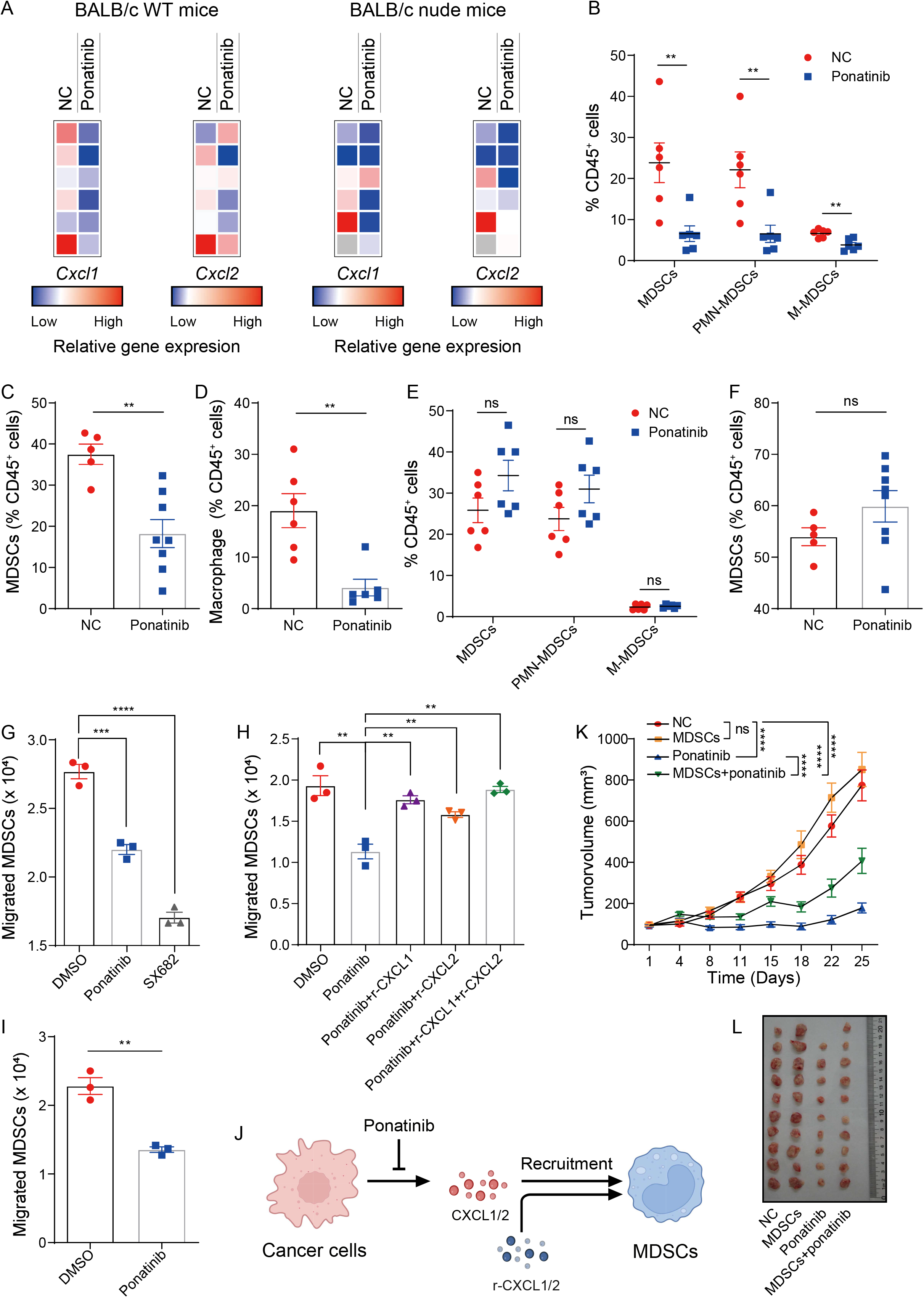
Ponatinib inhibits the infiltration of MDSCs into TME. (A) RT-qPCR analysis of *Cxcl1* and *Cxcl2* mRNA in 4T1 tumors from BALB/c WT mice (left) or nude mice (right) receiving the ponatinib treatments or control as described in Fig. 4B. Heat map representing the relative expression of indicated chemokine genes, data were normalized to Gapdh. Colors indicate the differences in gene expression (high mRNA level, red; low mRNA level, blue). Each square represents individual tumors from a single mouse (Grey square means not available, the indicated mouse sacrificed due to the requirement of animal ethics). (B) Quantification of flow cytometry results of tumor-infiltrating immune (CD45^+^) cells in 4T1 tumors from BALB/c WT mice treating with ponatinib or vehicle. Cell populations were identified as MDSCs (CD45^+^CD11b^+^F4/80^-^Gr-1^+^), PMN-MDSCs (CD45^+^CD11b^+^Ly-6G^+^Ly-6C^-^), and M-MDSCs (CD45^+^CD11b^+^Ly-6G^-^Ly-6C^+^). (C) Quantification of flow cytometry results of MDSCs in 4T1 tumors from BALB/c nude mice treating with ponatinib or vehicle. (D) Quantification of flow cytometry results of macrophage (CD45^+^ F80^+^) in 4T1 tumors from BALB/c WT mice treating with ponatinib or vehicle. (E) Quantification of flow cytometry results of MDSCs, PMN-MDSCs, and M-MDSCs cells in 4T1 spleens from BALB/c WT mice treating with ponatinib or vehicle. (F) Quantification of flow cytometry results of MDSCs in 4T1 spleens from BALB/c nude mice treating with ponatinib or vehicle. (G) Migration of MDSCs toward conditioned medium from 4T1 cells (ponatinib versus DMSO) were evaluated using *in vitro* transwell migration assay in triplicate. (H) Migration of MDSCs toward conditioned medium from 4T1 cells treated with DMSO or ponatinib supplemented with vehicle or recombinant mouse CXCL1 and CXCL2. Data represent mean ±SD. Statistical significance was determined by unpaired two-tailed Student’s t tests. \*\**P* < 0.01,\*\*\**P* < 0.001, \*\*\*\**P* < 0.0001. (I) Migration of MDSCs toward conditioned medium from MC38 cells (ponatinib versus DMSO) were evaluated using in vitro transwell migration assay in triplicate. (J) Illustration of ponatinib inhibiting CXCL1/CXCL2-MDSCs chemotaxis. (K) The mixture of 4T1 cells and MDSCs (isolated from spleen of tumor-bearing mice) were inoculated into the fat pad of BALB/c WT mice (n=10), and the growth of tumor was monitored. Ponatinib (30mg/kg) or vehicle was administered four days one week when tumor volume reached 100 mm³. Statistical significance was determined by two-way ANOVA. ns, *P* > 0.05, \*\*\*\**P* < 0.0001. (L) Images of 4T1 tumors of the indicated groups in (E). Data represent mean ±SEM. Statistical significance was determined by unpaired two-tailed Student’s t tests. \*\**P* < 0.01,\*\*\**P* < 0.001.

Thus, to assess the effect of ponatinib on tumor microenvironment, MDSCs and tumor-inhibiting leukocytes were analyzed in immune-competent tumor mouse models. We performed flow cytometry analysis of control-and ponatinib-treated tumors in WT mice to examine the distribution and trafficking of MDSCs. Indeed, our results showed that the proportion of MDSCs in 4T1 tumors were significantly reduced by ponatinib treatment (Fig.3B). In consistent with WT mice, we also observed a significant decrease of MDSCs infiltration in 4T1 tumors from BALB/c nude mice (Fig. 3C).

Next, we analyzed the subtype of MDSCs infiltrated in 4T1 tumors of WT mice after ponatinib treatment. It has been reported that MDSCs are subdivided into polymorphonuclear (PMN-) MDSCs and monocytic (M-) MDSCs (Youn et al., 2008). Upon ponatinib treatment, our results showed that both PMN-MDSCs and M-MDSCs were significantly decreased in the tumors (Fig. 3B). In addition, the infiltration of macrophages was also markedly decreased in the ponatinib-treated tumors (Fig. 3D).

To determine whether the decrease in MDSCs in the TME is due to infiltration process or a decrease in the overall number of MDSCs in the immune system, we also analyzed the MDSC population in the mouse spleen. Our results showed that there were no significant differences in the proportion of MDSCs in the spleens of either wild-type mice (Fig. 3E) or BALB/c nude mice (Fig. 3F) upon treatment with ponatinib, when compared to the control group. These results clearly demonstrate that the decreased presence of MDSCs in the TME after ponatinib treatment is a result of inhibited infiltration, not a decrease in the total number of MDSCs in the immune system.

To investigate whether the decrease in MDSCs in the TME was caused by the inhibited expression of CXCL1 and CXCL2 due to ponatinib treatment, a migration assay for MDSCs was conducted. We cultured freshly separated MDSCs from 4T1 tumors into the upper chamber of transwells, added the 4T1 condition media to the lower chamber, and then analyzed the effect of ponatinib on MDSCs migration ability.

As anticipated, treatment of 4T1 cells with ponatinib *in vitro* resulted in a significantly reduced migration of MDSCs into the growth medium (Fig. 3G). In addition, the receptor CXCR2, which plays a crucial role in MDSC trafficking into the tumor microenvironment (Katoh et al., 2013), was also analyzed. Our results showed that the addition of the CXCR2 inhibitor SX-682 to the growth medium derived from ponatinib treated 4T1 cells significantly reduced the migration of MDSCs (Fig. 3G). Furthermore, the involvement of CXCL1 or CXCL2 in MDSC recruitment under ponatinib treatment was confirmed by an increased migration of MDSCs upon the addition of recombinant mouse CXCL1 or CXCL2 protein to the growth medium of ponatinib treated 4T1 cells (Fig. 3H).

Notably, the similar results also observed in colon cancers. Our results showed that ponatinib treatment of MC38 cells significantly decreased the migration of MDSCs towards the growth medium *in vitro* (Fig. 3I). Together, our data suggested the infiltration of immunosuppressive MDSCs into TME is significantly inhibited by ponatinib treatment through suppressing the expression of CXCL1 and CXCL2 in cancer cells (Fig. 3J).

### The TNBC inhibition of ponatinib depends on the infiltration of MDSCs *in vivo*

To verify that the anti-tumor effect of ponatinib is gained through reducing the immune suppression of MDSCs, we conducted a MDSCs rescue experiment. MDSCs that isolated from tumor-bearing mice were mixed with 4T1 cells and injected into mammary pads of BALB/c mice. Our data showed that higher intratumoral MDSCs did not promote tumor growth without ponatinib treatment (Fig. 3K-3L). However, the growth of tumors was remarkedly accelerated in the presence of MDSCs mixture under ponatinib treatment (Fig. 3K-3L), confirming that anti-tumor effect of ponatinib was dependent on MDSCs. Therefore, the accumulation of MDSCs was compromised by ponatinib treatment, resulting in a moderation of tumor growth. Thus, reduction of immune-suppressive MDSCs is essential for the anti-tumor effect of ponatinib.

### Ponatinib enhanced the infiltration of “hot” tumor related immune cells into TME *in vivo* and exhibits no significant inhibition on immune cells in spleen

In addition to MDSCs, we also examined “hot” tumor related immune cells in TME after ponatinib treatment. Our results from WT mice showed that ponatinib caused significantly increases in the infiltration of CD3^+^ T cells to 4T1 tumor sites in contrast to control (Fig. 4A). Meanwhile, both CD4^+^ and CD8^+^ T cells are also greatly increased in the ponatinib-treated 4T1 tumors (Fig. 4A), implying that ponatinib promotes anti-tumor immune response in the mammary tumors.

**Fig4.**
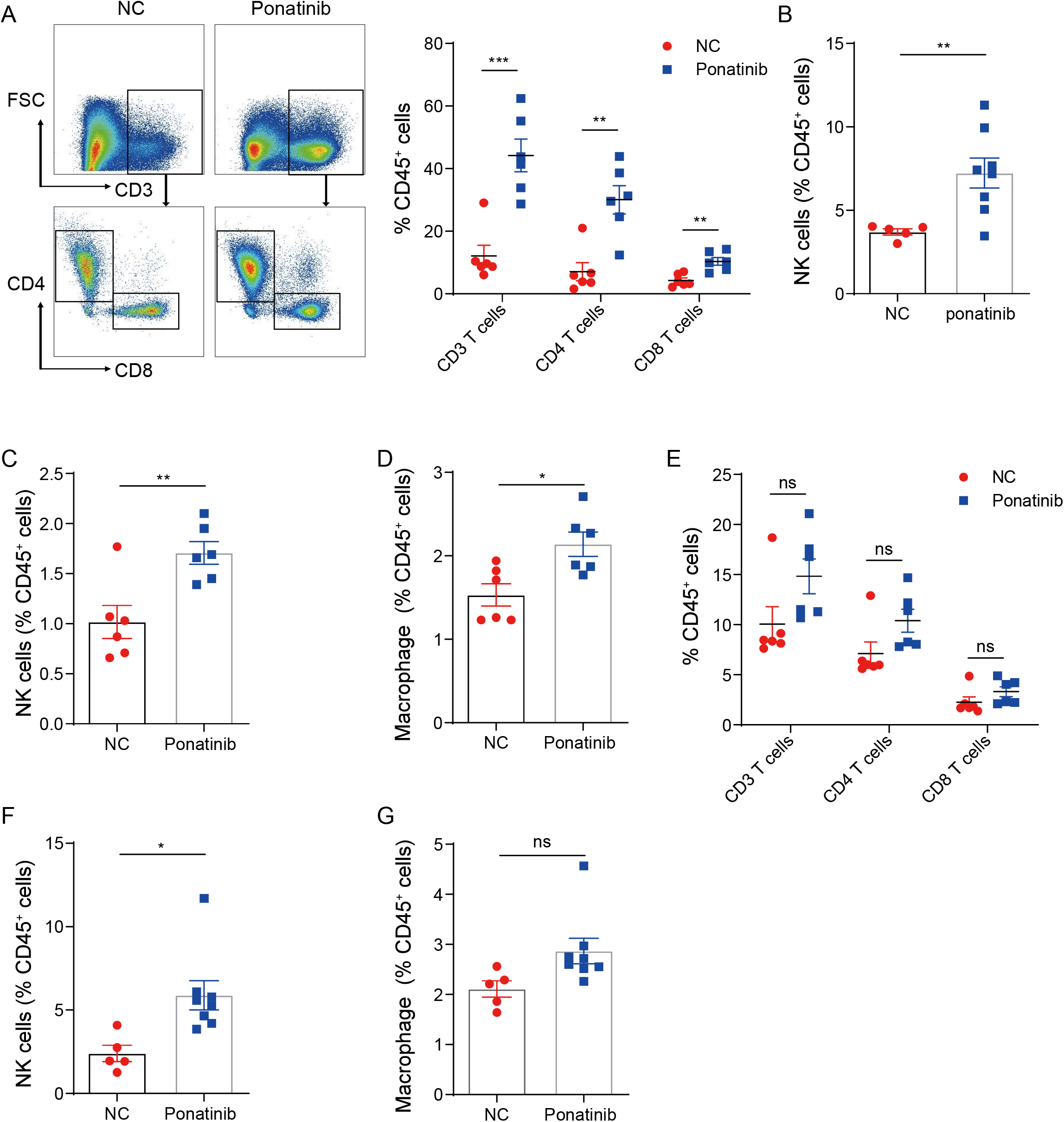
Ponatinib enhances the infiltration of “hot” tumor related T cells and NK cells into TME and exhibits no significant inhibition on immune cells in spleen. (A) Quantification of flow cytometry results of CD3^+^ T cells (CD45^+^CD3e^+^), CD4^+^ T cells (CD45^+^CD3e^+^CD8^-^CD4^+^) and CD8^+^ T cells (CD45^+^CD3e^+^CD8^+^CD4^-^) in 4T1 tumors rom BALB/c WT mice treating with ponatinib or vehicle as described in Fig.2A. (B) Quantification of flow cytometry results of NK cells (CD45^+^CD49b^+^) in 4T1 tumors from BALB/c nude mice treating with ponatinib or vehicle. (C to E)Quantification of flow cytometry results of (C) NK cells (CD45^+^Nkp46^+^), (D) macrophage and (E) CD3^+^ T cells, CD4^+^ T cells and CD8^+^ T cells in 4T1 spleens from BALB/c WT mice treating with ponatinib or vehicle. (F-G) Quantification of flow cytometry results of (F) NK cells (CD45^+^CD49b^+^) and (G) macrophage in 4T1 spleens from BALB/c nude mice treating with ponatinib or vehicle. Data represent mean ±SEM. Statistical significance was determined by unpaired two-tailed Student’s t tests. \*\**P* < 0.01,\*\*\**P* < 0.001.

Beside WT mice, we also examined the effect of ponatinib treatment on immune cells in 4T1 tumors from BALB/c nude mice. Even though there are no T cells in nude mice, notably, the proportion of NK cells in 4T1 tumors from BALB/c nude mice was remarkedly increased by ponatinib treatment (Fig.4B). These results indicate that ponatinib-induced NK cells enrichment in TME was not dependent on T cells.

In addition to examine immune cells in tumors, we also checked the effect of ponatinib on immune cells in mouse spleens. We observed higher percentages of NK cells (Fig. 4C) and macrophage (Fig. 4D) in spleens from WT mice under the treatment of ponatinib compared with control, while the other immune cell populations, including CD3 T cells, CD4 T cells and CD8 T cells, exhibited no significant differences (Fig. 4E). Furthermore, in line with the observations in spleens of the WT mice, ponatinib caused a dramatical increase of NK cells (Fig. 4F), but there were no significant differences in the distribution of macrophage in spleens of BALB/c nude mice under the treatment of ponatinib compared with control (Fig. 4G).

Collectively, our data suggested that ponatinib treatment leaded to obvious elevation of multiple types of “hot” tumor related T cells and NK cells in the microenvironment of the mammary tumor. This result is consistent with our finding that ponatinib treatment inhibited the infiltration of MDSCs into TME, as it is well known that MDSCs have suppressive effects on both T cells and NK cells proliferation and function. The decreased immunosuppressive MDSCs and increased “hot” tumor related T cells and NK cells in TME, induced by ponatinib treatment, at least partially, reprogrammed the “cold” immune microenvironment to “hot” one. Meanwhile, ponatinib treatment showed no significant inhibition on immune cells in mouse spleens, which indicating that the inhibition effect is specific to TME and not immune systemic.

### Ponatinib enhances the efficacy of immune checkpoint blockade therapy on TNBC *in vivo*

Given that ponatinib inhibited MDSCs infiltration into TME and thus enhanced T-lymphocyte responses, resulting a “hot” tumor microenvironment. We wondered whether ponatinib treatment might further enhance the efficacy of ICB. To evaluate our hypothesis, we performed *in vivo* studies with a syngeneic tumor model in which mice bearing 4T1 tumors were treated with ponatinib alone or combination with anti-PD-L1 antibody (Fig.5A). Our results show that ponatinib or anti-PD-L1 monotherapy did suppress tumor growth (56.7% and 38.5% inhibition ratio, respectively), however, there was a more pronounced tumor inhibition (74.6% inhibition ratio) with the combination of ponatinib and anti-PD-L1 over either single agent treatment (Fig.5B and 5C). Moreover, the survival curve results further confirmed that ponatinib significantly improves the survival of 4T1 tumor-bearing mice with anti-PD-L1 treatment (Fig.5D). Together, these results demonstrated that ponatinib treatment could sensitize the malignant breast cancer to checkpoint inhibitor-based immunotherapy through reshaping the tumor microenvironment from “cold” tumors to “hot” tumors.

**Fig5.**
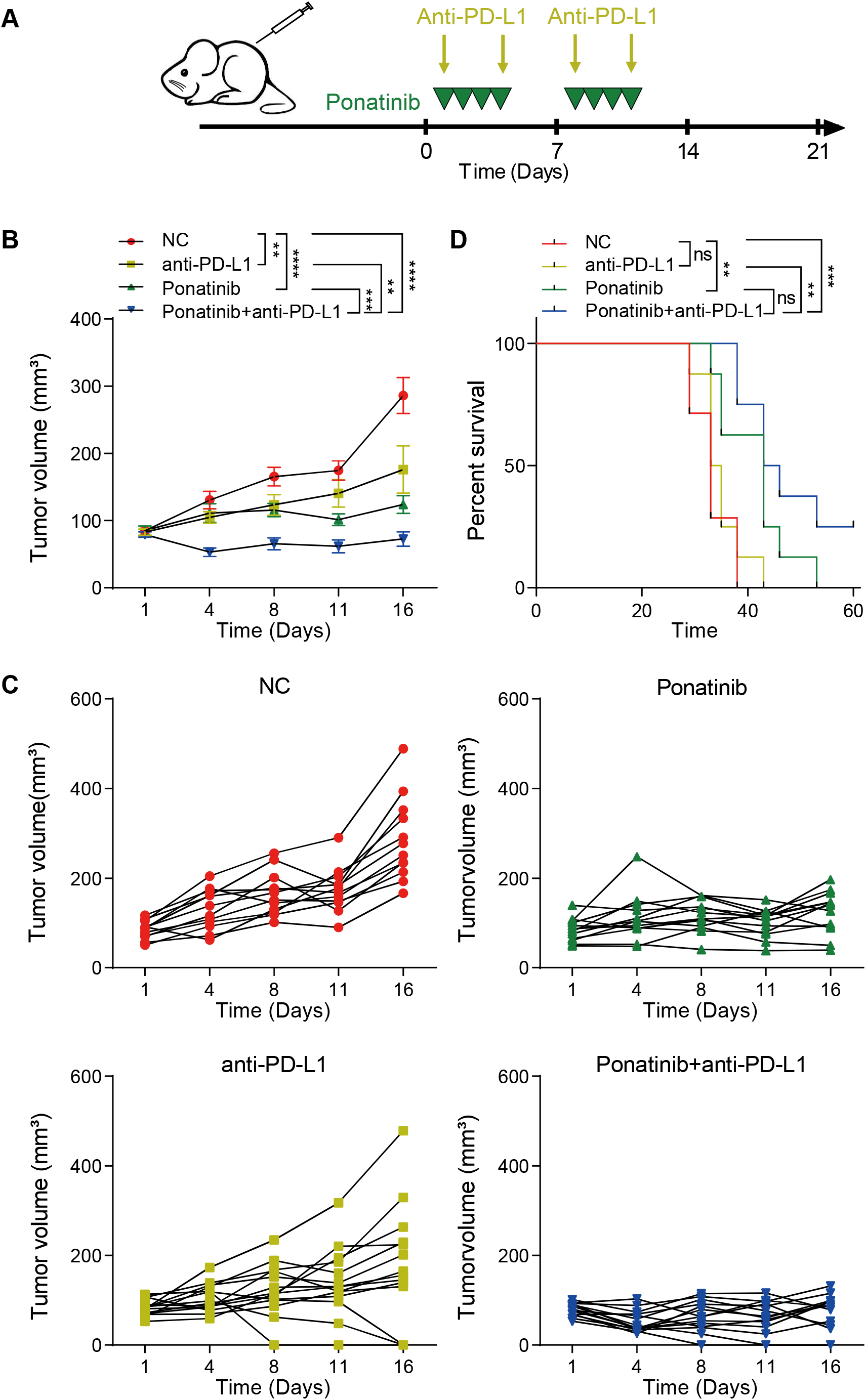
Combination treatment with ponatinib enhances the therapeutic efficacy of PD-L1 in TNBC. (A) Schematic diagram for the in vivo combination treatment in 4T1-bearing mice. Ponatinib (30mg/kg) or vehicle was administered four days one week and PD-L1 mAb (200 μg/injection, twice a week) alone or in combination (n=12-14 mice/group) when tumor volume reached 100 mm³. (B and C) Growth of 4T1 cells in BALB/c WT mice treated with the indicated conditions in (A). Data represent mean ± SEM of n mice, as indicated on each panel. Statistical significance was determined by two-way ANOVA. \*\**P* < 0.01, \*\*\**P* < 0.001, \*\*\*\**P* < 0.0001. (D) Survival curves for each treatment group. Statistical significance was determined by log-rank (Mantel-Cox) test. ns, *P* > 0.05, \*\**P* < 0.01 and \*\*\**P* < 0.001.

### p38 induced STAT1 phosphorylation mediates the inhibition of CXCL1 and CXCL2 expression by ponatinib

To further understand how ponatinib regulates the expression of “cold”-tumor related genes, we examined whether ponatinib suppresses the expression of CXCL1 and CXCL2 through its well-known targets SRC and ABL. Our results showed that knocking down the expression of *SRC*, *ABL1* or *ABL2* by shRNAs in cancer cells exhibited no significant difference on the mRNA level of *CXCL1* and *CXCL2* (Fig. S1A and Fig. S1B), which indicated that SRC or ABL might not be the targets of ponatinib for the inhibition of CXCL1 and CXCL2 gene expression.

Then, we conducted a follow-up RNA sequencing (RNA-seq) study to examine transcriptome-wide effects of ponatinib with both MDA-MB-231 cells and 4T1 cells. The gene set enrichment analysis (GSEA) showed that MAPK signaling pathways were significantly enriched (Fig. 6A). To investigate how MAPK pathway activation involved with response to ponatinib treatment, we evaluated the phosphorylation levels of ERK, p38 and JNK, as they are the three main molecules downstream of the MAPK pathway. Western blot analysis showed that only the phosphorylation of p38 was significantly decreased in MDA-MB-231 cells (Fig. 6B) and other cancer cell lines (Fig. S2A-2E) treated with ponatinib.

Previous work reported that the phosphorylation level of STAT1 Ser727 was closely related to the p38 MAPK signaling pathway(Goh et al., 1999). Our western blot analysis also revealed that knocking down p38 significantly suppressed the phosphorylation of STAT1 at Ser727 (Fig.6C). In line with the observations of knocking down of p38, western blot analysis showed a significant decrease phosphorylation level of STAT1 at Ser727, but not Tyr701, in MDA-MB-231 cells and 4T1 cells (Fig. 6D) and multiple other cancer cells (Fig.S2C and Fig.S2D). In addition, we also revealed that there was no significant change in the phosphorylation level of STAT1 after knocking down SRC or ABL in MDA-MB-231 and 4T1 cells (Fig. S2F-2G). Together, our results clearly showed that p38 phosphorylation and consequently STAT1 phosphorylation are significantly inhibited in cancer cells by ponatinib treatment.

**Fig6.**
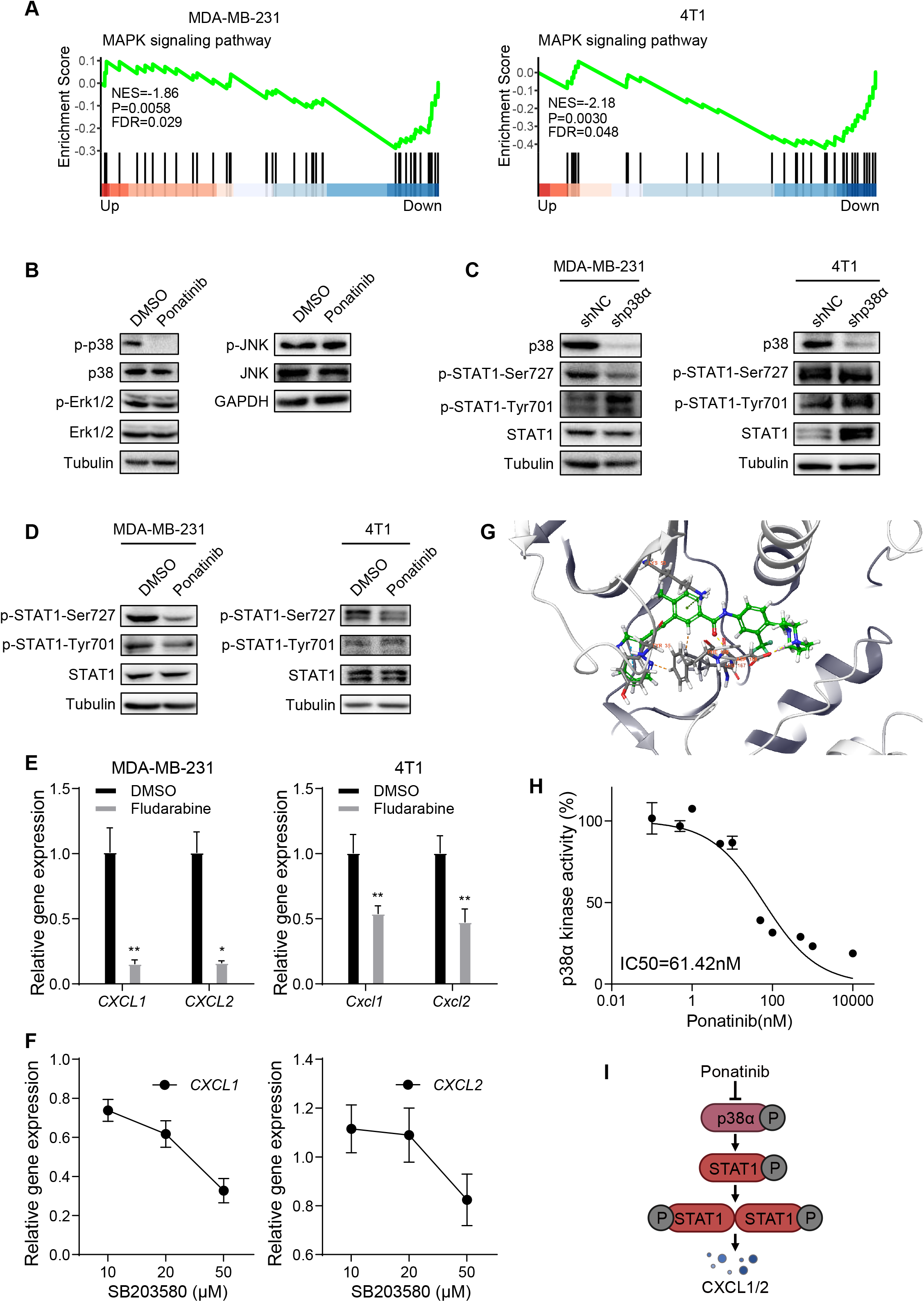
Ponatinib inhibits the expression of *CXCL1* and *CXCL2* through p38-STAT1 pathway. (A) GSEA analysis of MAPK signaling pathway in DMSO- or Ponatinib-treated cells (left, MDA-MB-231 cells; right, 4T1 cells). NES, normalized enrichment score. *P* value represents false discovery rate adjusted *P*-value. (B) Western blot analysis of the indicated protein in MDA-MB-231 cells treated with 1μM ponatinib or DMSO. (C) Western blot analysis of the indicated protein in MDA-MB-231 cells (left) or 4T1 cells (right) knocking down p38α compared with negative control. (D) Western blot analysis of the indicated protein in MDA-MB-231 cells (left) or 4T1 cells (right) treated with 1μM ponatinib or DMSO. (E) RT-qPCR analysis of *CXCL1* and *CXCL2* mRNA in in MDA-MB-231 cells (left) or 4T1 cells (right) cells treated with fludarabine or DMSO. (F) RT-qPCR analysis of *CXCL1* and *CXCL2* mRNA in in MDA-MB-231 cells treated with different concentrations of SB203580 at 24 h. Data represent mean ± SD of three independent experiments. Statistical significance was calculated by unpaired two-tailed Student’s t test. \**P*< 0.05 and \*\**P*< 0.01. (G) 3D interaction diagram of ponatinib in the active site of p38α. (H) *In vitro* p38α activity treated with different concentrations of ponatinib. Data represent mean ± SD of three independent experiments and normalized to DMSO control. (I) Illustration of ponatinib inhibiting CXCL1/CXCL2 expression through inhibiting p38-STAT1 pathway.

Next, we aimed to study whether the inhibition of ponatinib on CXCL1 and CXCL2 is carried out through p38 mediated STAT1 phosphorylation. Previous work has shown that the promoter regions of *CXCL1* and *CXCL2* have multiple potential binding sites for STAT1 (Burke et al., 2014). Our data showed that the mRNA level of *CXCL1* was significantly decreased after knocking down STAT1, and the expression of *CXCL2* was also shown the similar trend (Fig. S3A-S3B). To further validate the regulation of STAT1 on the expression of *CXCL1* and *CXCL2*, we treated cancer cells with STAT1 inhibitor fludarabine. Our results showed that the expression of *CXCL1* and *CXCL2* were also significantly inhibited in MDA-MB-231 cells and 4T1 cells (Fig. 6E). In addition, downregulation of *CXCL1* and *CXCL2* were further confirmed in cancer cells treated with p38 inhibitor SB203580 (Fig. 6F). Collectively, these results indicated that ponatinib inhibits the expression of *CXCL1* and *CXCL2* might through p38-STAT1 signaling pathway.

### Ponatinib directly inhibits kinase activity of p38α

Given that ponatinib decreased the phosphorylation level of p38 in various cancer cell lines, we wondered whether ponatinib could directly bind to and affect its kinase activity. The molecular docking results revealed significantly high binding affinity between ponatinib and p38α among the indicated proteins according to the lowest docking score, which refers to free energy (Fig.6G and TableS1). Most importantly, our results from the p38α kinase assay clearly demonstrated that the kinase activity is significantly inhibited by ponatinib in a dose-dependent manner (Fig.6H). Together, these results suggested that ponatinib might directly bind to p38α and inhibit its phosphorylation and kinase activity, which in turn reduces the phosphorylation of STAT1 and thus the expression of CXCL1 and CXCL2 (Fig. 6I).

## Discussion

In this study, we identified ponatinib as the ICB synergetic agent through a “cold”- tumor related gene signature-based high throughput screening. Particularly, in proof-of-concept studies, we establish that therapeutic inhibition of MDSC recruitment through targeting p38-STAT1 signaling improves anti-tumor immune response and sensitivity to ICB therapy for TNBC (Fig.7).

**Fig7.**
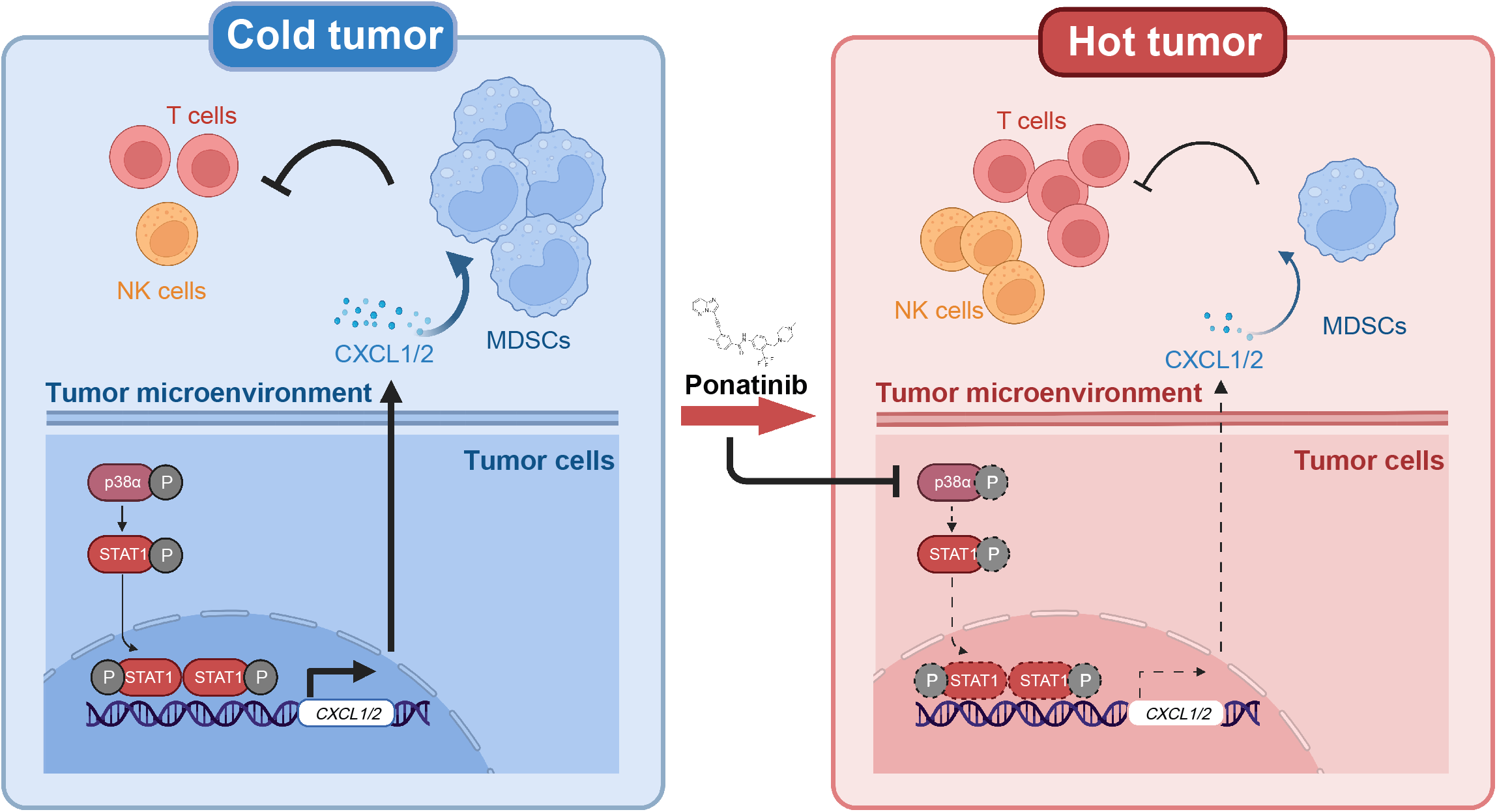
Schematic illustration of the mechanism of ponatinib inhibiting CXCL1/CXCL2 expression and MDSCs recruitment through inhibition of p38α-STAT1 pathway.

MDSCs mediated immune suppression is a major mechanism responsible for immune evasion in tumors. Moreover, MDSC is considered as a major source of drug resistance in anti–PD-1/PD-L1 immunotherapy (Weber et al., 2018). In this study, we demonstrated that in tumor-bearing mice, ponatinib directly modulates the immune microenvironment of breast tumors via inhibition of tumor-promoting MDSCs and restoration of antitumor T cells or NK cells, resulting in inhibition of tumor growth (Fig.7). Our findings emphasize that ponatinib inhibit the immunosuppressive TME via the CXCL1/2-CXCR2 pathway and MDSCs, supporting to combination therapeutic strategies for TNBC.

This new mechanism of action might contribute to the understanding of the efficacy of ponatinib in cancers. Ponatinib is an FDA-approved drug for leukemia, it is unclear whether and how this compound directly modulates the immune microenvironment. As 4T1 tumor model is a mesenchymal-like triple-negative cancer model, the obvious therapeutic effect that ponatinib sensitizes 4T1 tumors to anti–PD- L1 therapy (Fig.7), suggesting that this small molecule could convert the relative "cold" tumors to "hot". Currently, various clinical trials which combined ICIs and MDSCs targeting have been applied in different tumor types. Together, our results suggest new chemoimmunotherapeutic approaches combining ponatinib with PD-1/PD-L1 antibodies could be effective strategies in treatment of human advanced breast cancer including triple-negative subtype.

Our investigations of the molecular signaling pathways associated with ponatinib treatment revealed that the inhibition of “cold”-tumor related chemokines *CXCL1* and *CXCL2* involves p38α and STAT1. These discoveries are consistent with previous studies which reported that p38α inhibition can significantly decrease STAT1 phosphorylation and inhibit its function (Goh et al., 1999) and reports that STAT1may bind to the promoter of *CXCL1* and *CXCL2* (Burke et al., 2014). Moreover, our results demonstrated that ponatinib could directly inhibit the kinase activity of p38α, which is consistent with previous report (Worzella et al., 2017). In addition to p38α, our results indicate that STAT1 might be another promising target for the combinational immunotherapy and need to be studied further.

Our study emphasizes that ponatinib can serve to systematically reprogram the immunosuppressive TME to enhance the T cell responses via the p38α-STAT1- CXCL1/2 pathway and immune-suppressive myeloid cells. Our findings indicate a complex effect of ponatinib on both cancer cells and the tumor immune cell environment. A clear understanding of these signaling interactions might enable the design of new therapeutic strategies that exploit the highly impactful role of immunosuppressive cells infiltration on the efficacy of immunotherapy. In the combination treatment experiment, ponatinib was administrated for 2 weeks and PD- L1 was administrated 4 times in this study. It was shown that the progression of the tumors was significantly delayed. However, long-term efficacy and the generation of resistance for ponatinib in breast cancer need to be assessed in the future.

In summary, our study provides the possibility for developing new approaches to modulate TME for enhanced anti-tumor immunity, discovering the anti-”cold”-tumor-microenvironment drugs as a combination agent for immunotherapy with anti-PD-L1. Applying this new strategy, we identify and demonstrate that the chemical inhibition of P38α by ponatinib could decrease the abundance of CXCL1 and CXCL2, block the infiltration of MDSCs into tumor microenvironment and thus enhance the efficacy of immunotherapy in TNBC.

## Materials and methods

### Study design

We used our HTS^2^ approach (Li et al., 2012; Shao et al., 2019) to screen 1,154 FDA approved by monitoring their effects on the expression of *CXCL1* and *CXCL2*. RT-qPCR and western blot analysis were used for investigation of candidate compounds. All of the *in vitro* and *in vivo* studies were performed using the breast cancer and colorectal cancer cell lines, or Balb/c mice, Balb/c nude mice, NSG mice and C57BL/6 mice to detect candidate compounds that inhibit MDSCs infiltration into the tumor microenvironment and that improve anti-PD-L1 efficacy in controlling tumor growth.

### Chemicals

The screening library (1,154 FDA approved drugs) were purchased from Selleck. Ponatinib were purchased from Targetmol. All chemicals were dissolved in dimethyl sulfoxide (DMSO) for the *in vitro* studies.

### Drug screening and enrich score calculation

The drug screening data used in this project was generated by Shao, et al (Shao et al., 2019). The methods of calculating the activity score to reverse "cold" tumor into "hot" tumor for each compound was similar with method described in the original connectivity map literature (Lamb et al., 2006) and in our previous work (Lin et al., 2020; Shao et al., 2019; Wang et al., 2021).

4 genes (those expressed in MDA-MB-231 cell lines and with available screening probes) including *CXCL10*, *CXCL11*, *CXCL1* and *CXCL2* were combined to serve as the "query" gene set, and the HTS^2^ expression profiles of all screened compounds were served as the "target" gene sets, and Kolmogorov-Smirnov test was used to compare the similarity of each paired "query" and "target" to assign a final similarity score for each compound.

### Cell culture

The MDA-MB-231, 4T1, LM2, MCF7, SW480, SW620, HCT116, Hela and A549 cell lines were obtained from the Institute Cancer Cell Line Encyclopedia of China. MC38 and B16F10 cell lines were kindly provided by Dr. Deng Pan from Tsinghua University. Cells were cultured in DMEM (Gibco) supplemented with 10% fetal bovine serum (Gemini) and 1% penicillin/streptomycin (Gibco). Cells were cultured at 37 °C with a 5% CO_2_ atmosphere.

### Plasmids and antibodies

Lentiviral shRNAs were provided by the Vector Core at Tsinghua University. Antibodies used included: anti-p38, anti-p-p38-Thr180/Tyr182, anti-stat1, anti-p- STAT1-Tyr701, anti-p-STAT1-Ser727, anti-p-Erk1/2, anti-Erk1/2, anti-p-JNK, anti-JNK, anti-GAPDH, anti-tubulin were purchased from Cell Signaling Technology.

### Mice and reagents

The laboratory animal facility has been accredited by AAALAC (Association for Assessment and Accreditation of Laboratory Animal Care International) and the IACUC (Institutional Animal Care and Use Committee) of Tsinghua University approved all animal protocols used in this study.

Female mice (5–6 weeks old) were housed in the animal facility of the Laboratory Animal Research Center of Tsinghua University under specific pathogen-free conditions. Briefly, sub confluent 4T1 cells (5×10 ^4^) were suspended in 0.04 ml DMEM medium and injected into the mammary fat pads of mice (BALB/c WT mice, BALB/c nude mice and NSG mice) under anesthesia with avertin. For MC38 tumor model, C57BL/6 female mice were subcutaneously injected in the right flank with tumor cells (1 × 10^6^ cells). Animals were administrated when the volumes of tumors were about 100 mm^3^. Mice were treated with ponatinib at 30 mg/kg or vehicle orally fourth weekly for 3 weeks.

For combination therapy, ponatinib treatment was administered for 2 weeks combining with an i.p. injection of either a-PD-L1 antibody (200μg twice weekly for 2 weeks, a total of 4 times; clone 10F.9G2, BioXcell) or isotype control antibody (200μg twice weekly for 2 weeks, a total of 4 times; clone LTF2, BioXcell).

Tumor volumes were measured with digital calipers and were calculated with the formula: volume (mm^3^) = [width^2^ (mm^2^) × length (mm)]/2. After 22 days from the beginning of treatment, mice were sacrificed. Tumors were then isolated for FACS.

### MDSCs mixed with 4T1 cells for transplantation

4T1 cells (5×10 ^4^) with sorted MDSCs (1×10 ^5^) from 4T1 tumor-bearing mice were injected into the mammary gland. The tumor growth curves and tumor weight were recorded.

### MDSC isolation and *in vitro* migration assay

MDSCs were isolated from the spleens of 4T1 tumor-bearing mice using a Mouse MDSC Isolation Kit (Miltenyi Biotec, Cat# 130-094-538) and plated in RPMI1640 supplemented with 10% FBS and antibiotics. MDSCs (1×10 ^5^ cells/well) were seeded in the top chamber of the transwell (Corning). Conditioned media (CM) from cultured 4T1 cells or MC38 cells treating with 1μmol/L ponatinib or DMSO for 48 hours were collected, supplemented with vehicle or recombinant mouse CXCL1 and CXCL2, and added to the bottom layer of the transwell. After 6 hr incubation, cells that had completely migrated to the bottom chamber were counted. These experiments were performed in triplicate, and statistical significance was assessed using Student’s t-test.

### Flow cytometry analysis

For flow cytometry, splenocytes, and tumor infiltrating lymphocytes were isolated, and red blood cells were lysed. The single cell samples were incubated with anti-mouse CD16/32 (BioLegend) for 10mins at 4℃. Then the single-cell suspensions were stained with indicated antibodies for 40 min on ice: anti-mouse CD4 Super Bright 600(eBioscience), anti-mouse CD8a PE-eFluor 610 (eBioscience),anti-mouse CD3e PerCP-Cyanine 5.5 (eBioscience), anti-mouse CD3e PE-Cyanine 5.5 (eBioscience), anti-mouse CD45 eFluor 450(eBioscience), anti-mouse CD45 Super Bright 780 (eBioscience), anti-mouse Ly-6G/Ly-6C PE- Cyanine7 (eBioscience), anti-mouse Ly- 6G-PE(eBioscience), anti-mouse Ly-6C eFluor 450 (eBioscience), anti-mouse F4/80 PE-CY7 (eBioscience), anti-mouse F4/80 FITC (eBioscience), anti-mouse CD11b fluoresceinisothiocyanate (FITC) (eBioscience), anti-mouse CD11b eFluor 506 (FITC) (eBioscience). Flow cytometry was performed using flow cytometer FACS AriaIII (BD) and analyzed with FlowJo software.

### RT-qPCR

RNA that was exacted with TRIzol (Invitrogen) was used for cDNA synthesis with a Thermo Scientific RevertAid First Strand cDNA Synthesis Kit (ThermoFisher Scientific). RT-qPCR was performed using a StepOne Plus real-time PCR system (KAPA).

### ELISA

Cells (2×10 ^6^/100 mm dish) were cultured for 24 hr. Media were removed and replaced with 10 ml fresh DMEM containing different concentrations of ponatinib or DMSO. Supernatants were collected 24hr or 48 hr later with any floating cells removed by 0.45 mm filtration. The amount of CXCL1 or CXCL2 protein in the supernatant was determined using a mouse CXCL1 or CXCL2 specific ELISA kit (R&D system). All experiments were performed according to the manufacturer’s instructions.

### Immunoblotting

Cells were lysed with RIPA lysis buffer containing 1 mmol/L PMSF and 1× protease inhibitor cocktail (MedChemExpress). After BCA protein quantification (Tentler et al.), protein samples were separated by SDA-PAGE gel and transferred to polyvinylidene membranes (Bio-Rad), and probed with antibodies indicated in the legends of each relevant figure. The bands were visualized by using SuperSignal West Pico reagent (ThermoFisher Scientific). GAPDH or Tubulin was used as an internal control.

### RNA-sequencing (RNA-seq)

1×10 ^6^ MDA-MB-231 cells were plated in a 10cm-well plate for 24 h and then treated with ponatinib at 1 µmol/L for 24 h. 2 × 10 ^6^ 4T1 cells were plated in a 6-well plate for 24 h and then treated with ponatinib at 1 µmol/L for 24 h. Cells were harvested, and total RNA was isolated using TRIzol (Invitrogen). The libraries were constructed by using the NEBNext Ultra RNA Library Prep Kit for Illumina (NEB) and sequenced on NovaSeq6000 sequencing system (Illumina).

Clean reads were obtained after removing adapter, poly-N sequences. These reads were mapped to the reference genome Homo sapiens (hg38) and Mus musculus (GRCm39) (Howe et al., 2021) using HISAT2 tools (Kim et al., 2019), respectively. Then, gene read counts were obtained by assembling transcripts using HTSeq (Anders et al., 2015) for each sample. DESeq2 were used to perform differentially expressed analysis, and obtain differentially expressed genes (DEGs) with threshold of P value < 0.05 and fold change value ≥ 1.5. We used clusterProfiler package (Wu et al., 2021) to performed KEGG pathways enrichment and GSEA analysis for DEGs.

### Gene-set enrichment analysis (GSEA)

We used clusterProfiler package (Wu et al., 2021) to performed KEGG pathways enrichment and GSEA analysis for DEGs.

### Molecular docking

The crystal structures of p38 protein were obtained from the RCSB Protein Data Bank (PDB). Then, the crystal structure of each protein was selected based on the optimal available resolution. Moreover, the protein preparation wizard module of Schrodinger’s Maestro Molecular modeling suit (Schrödinger Release, 2019 –1) was utilized to prepare the protein crystallographic structures. In addition, the LigPrep module of Schrodinger’s Maestro Molecular modeling suit was employed to obtain the 3D structures and energy minimization of the active ingredients in GFD. Based on the specific known active sites of the protein targets, Glide was adopted for all molecular docking simulations and calculations. Moreover, the ligand-target interactions were visualized by the ligand interaction diagram module.

### p38α Kinase Assay

The p38α kinase assay was performed according to the manufacturer’s instructions. Reactions were performed in 25µl mixture containing 1 **×** Reaction Buffer A, 1% DMSO, 50µM DTT, 10 µM ATP, 40ng of p38α kinase, 1µg p38α peptide substrate and serial dilution of ponatinib. As a control for reagent resistance to the compounds, the same titration was performed with all the reaction components except the enzymes. The reactions were incubated at room temperature for 60 minutes. After the incubation time, 25µl of Kinase-Glo® Reagent was added to the reactions and the plate was incubated at room temperature for 40 minutes. Then Kinase Detection Reagent was added to the reactions and the plate was incubated at room temperature for 30 minutes. Luminescence was recorded and IC50 values were determined.

### Statistical analysis

Two-Way ANOVA and Two-tailed unpaired Student’s t-tests were used for comparisons the differences of two groups; Log-rank tests were used for Kaplan-Meier survival analysis. The level of significance was set at \**P* < 0.05, \*\**P* < 0.01, \*\*\**P* < 0.001 and\*\*\*\**P* < 0.0001.

### Code availability

All codes used in this study were followed the manuals of each R package, which are fully open accessed.

## Acknowledgments

We thank the staff members of the Animal Facility of the Laboratory Animal Research Center of Tsinghua University.

## Author contributions

D.W. and Q.W. jointly conceived the main conceptual ideas. Q.W. designed and carried out the experiments and performed data analyses. S.L and K.L provided supports in drug screening data analysis. Y.W. analyzed and processed sequencing data. Y.D. contributed to molecular docking. All authors coordinated the work. D.W., Q.W. and G.Z. wrote the manuscript with input from all the other authors.

## Data and materials availability

All data supporting our conclusion of this study are presented in the article. All other data relevant to this paper may be requested from the authors.

