## SUPPLEMENTARY INFORMATION for "Reversing myeloid-derived suppressor cells mediated immunosuppression via p38α inhibition enhances immunotherapy efficacy in triple negative breast cancer"

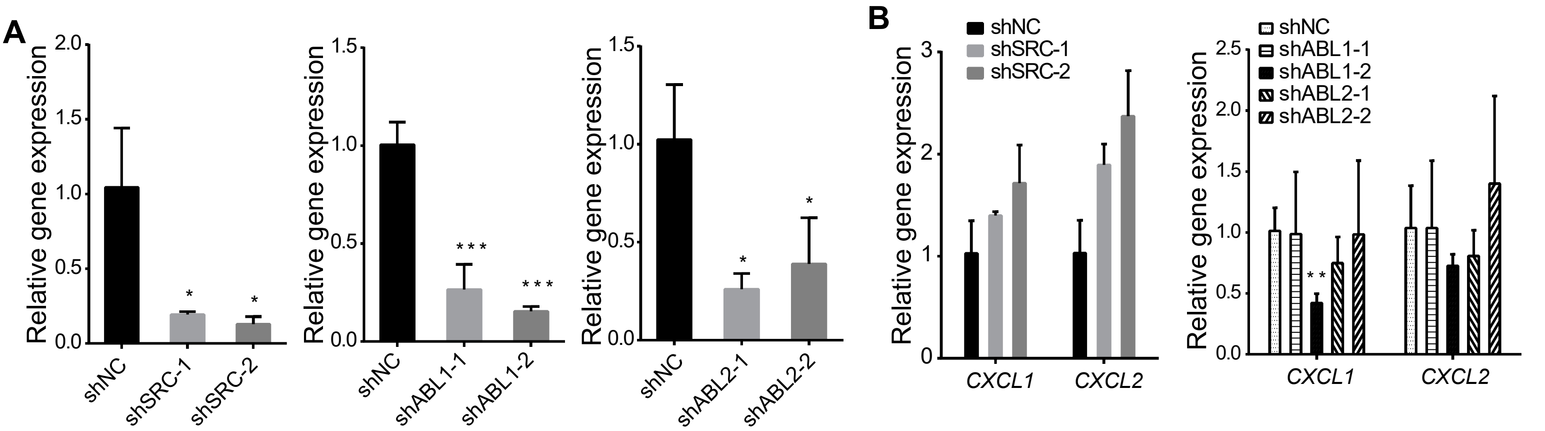


FigS1. SRC/ABL1/ABL2 are not the targets for the inhibition of CXCL1 and CXCL2 expression in TNBC cells by ponatinib. .

**FigS1. SRC/ABL1/ABL2 are not the targets for the inhibition of CXCL1 and CXCL2 expression in TNBC cells by ponatinib.**

(A and B) RT-qPCR analysis of (A) knockdown efficiency or (B) *CXCL1* and *CXCL2* mRNA in MDA-MB-231 cells. The RT-qPCR data were normalized to GAPDH.

The RT-qPCR data were normalized to GAPDH. Data represent mean ± SD of three independent experiments. Statistical significance was calculated by unpaired two-tailed Student's t test. *$P$< 0.05, **$P$< 0.01 and ****$P$< 0.001.

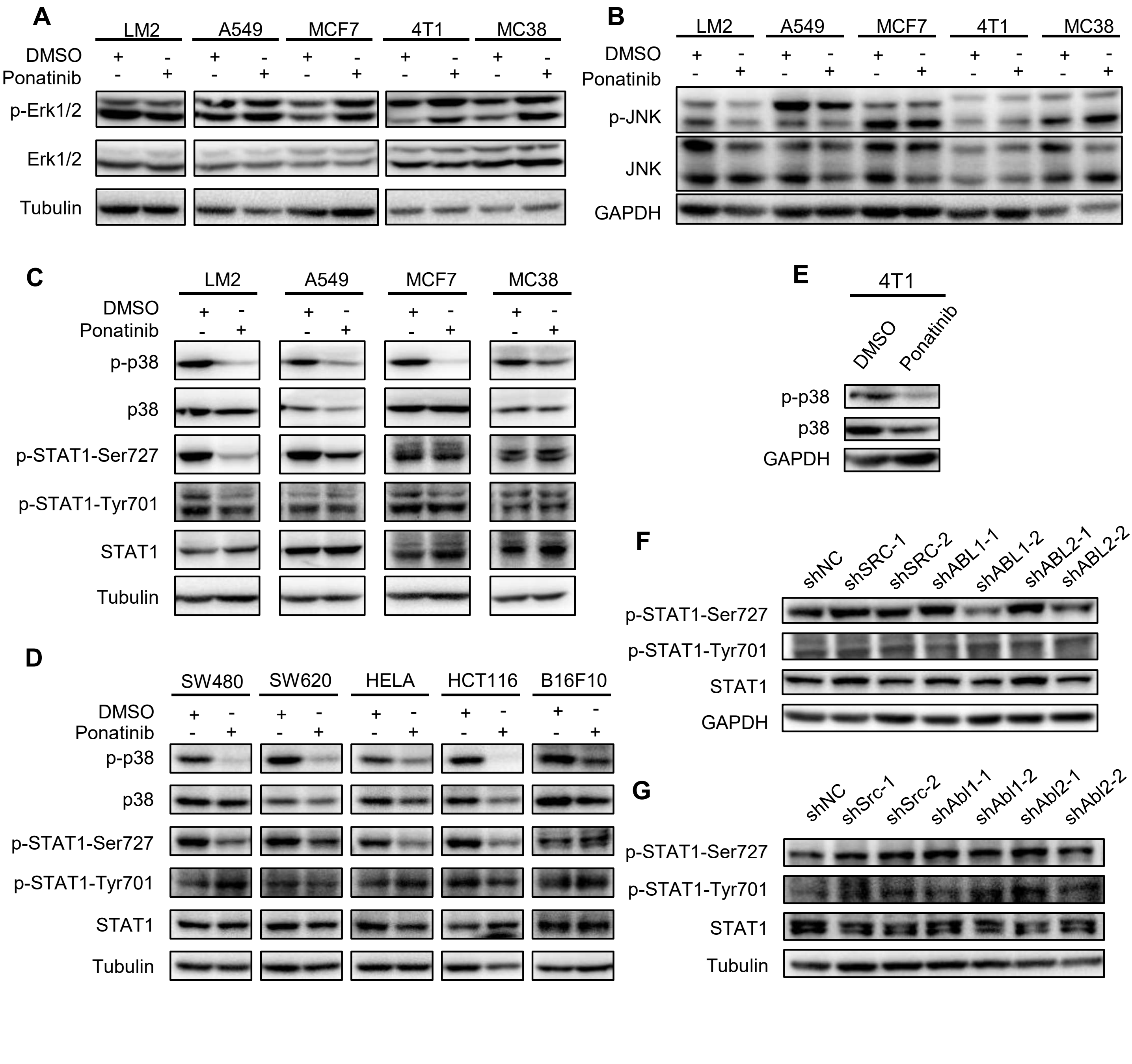


FigS2. Ponatinib inhibits p38-STAT1 pathway in multiple cancer cell lines.

**FigS2. Ponatinib inhibits p38-STAT1 pathway in multiple cancer cell lines.**

(A to E) Western blot analysis of the indicated protein in multiple cancer cell lines treated with 1μM ponatinib or DMSO.

(F and G) Western blot analysis of the indicated proteins in MDA-MB-231 cells (F) or 4T1 cells (G) knocking down SRC/ABL1/ABL2 compared with negative control.

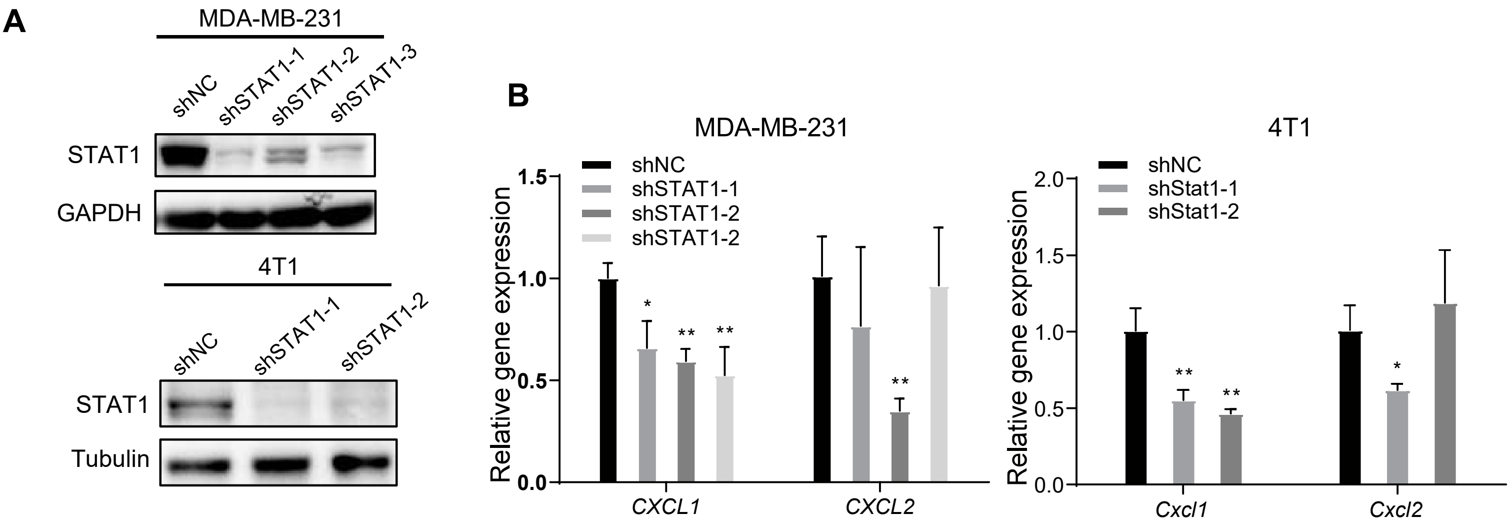


FigS3. STAT1 inhibits the expression of *CXCL1* and *CXCL2*.

.

**FigS3. STAT1 inhibits the expression of *CXCL1* and *CXCL2*.**

(A) Western blot analysis shows the knockdown efficiency of STAT1 in MDA-MB-231 cells (up) or 4T1 cells (down).

(B) RT-qPCR analysis of *CXCL1* and *CXCL2* mRNA in in MDA-MB-231 cells (left) or 4T1 cells (right) knocking down the indicated genes compared with negative control. The RT-qPCR data were normalized to GAPDH. Data represent mean ± SD of three independent experiments. Statistical significance was calculated by unpaired two-tailed Student's t test. *$P$< 0.05 and **$P$< 0.01.

**Table S1. The results of molecular docking.**

| Protein | docking score | XP GScore | MM-GBSA dG Bind |
|---|---|---|---|
| MAPK11 | -10.506 | -10.712 | -78.58 |
| MAPK12 | -3.772 | -3.979 | -52.83 |
| MAPK13 | -7.038 | -7.244 | -64.02 |
| MAPK14 (p38α) | -12.854 | -13.609 | -84.89 |
| MAP2K6 | -4.624 | -4.831 | -54.91 |
